# Internal representation of future interactions in rats

**DOI:** 10.1101/2023.12.22.573019

**Authors:** T Dvorakova, V Lobellova, P Manubens-Coda, A Sanchez-Jimenez, JA Villacorta-Atienza, A Stuchlik, D Levcik

## Abstract

Animals and humans receive the most critical information from parts of the environment that are immediately inaccessible, possibly only visually explored, and highly dynamic. The brain must effectively process potential interactions between elements in such an environment to make appropriate decisions in critical situations. We trained male Long-Evans rats to discriminate static and dynamic visuospatial stimuli displayed on an inaccessible computer screen. We demonstrated that the rats learned to discriminate dynamic visuospatial stimuli faster, with greater accuracy and shorter reaction time than complementary static stimuli. Furthermore, we provide behavioral evidence indicating that rats internally represent dynamic environments as static maps that capture meaningful future interactions. These observations highlight the ecological importance of dynamic stimuli in the outside world and support previous findings in humans that internal static representations can encapsulate relevant spatiotemporal information of dynamic environments. Such a mechanism would allow animals and humans to process complex time-changing situations neatly.

## INTRODUCTION

Spatial cognition is essential for most animal species, including humans, as it allows them to locate resources efficiently, avoid obstacles and predators, and navigate complex environments. This ability is allowed by the so-called cognitive map – a mental representation of the physical environment^1^. Such a map includes spatial relationships between objects and places and their relative distances and directions.

Static environments, where the locations of objects and landmarks remain constant, allow individuals to form stable cognitive maps based on their experiences and interactions with that environment. However, the real world is inherently unstable. Spatial cognition must operate in dynamic environments, where multiple objects may change locations with variable velocities and directions, demanding continuous real-time updating and coordination of information from different sources. Despite the abundant wealth of knowledge regarding spatial cognition, there remains a significant gap in understanding the processing of dynamic spatial information derived from sources beyond the immediate reach of the individual.

Animals must represent their position within an environment and the positions of other objects in their surroundings. Laboratory spatial tasks for rodents typically require navigation to target places within stable environments^2,3^ or approaching and contacting other static objects to discriminate their positions^4,5^. Therefore, in most experimental paradigms, items of interest are placed within accessible parts of a familiar environment so that animals can associate previously visited positions with objects’ locations^6^. However, in natural conditions, important information often comes from distal (visual) parts of the environment without direct access.

How rodents perceive visual stimuli became more widely studied with the advent of behavioral tasks that utilized computer screens and virtual environments. Such an experimental setup enables complete automation and precise control over presented stimuli and introduces analogous versions of human tasks for rodents. However, in these tasks assessing spatial memory^7^, paired-associate learning^8^, attention^9^, or cognitive flexibility^10^, visual stimuli are static and presented through touchscreens, and animals are trained to physically approach and make contact with the visual stimuli. Further tasks were designed to study the visual perception of motion^11,12^ or discrimination between self-motion and the motion of other objects^13^. Rats can also discriminate positions of visual objects, which cannot be directly explored and are displayed on a distant computer screen in an inaccessible space^14,15^ and this ability is dependent on the hippocampus^16^, which is critical for navigation in the real space, both in static^17^ and dynamic scenarios^18^. However, the particular process of encoding dynamic objects and their impending interactions beyond the accessible space have not yet been understood.

A mechanism called time compaction could underlie the processing of distal dynamic situations^19^. This mechanism introduces a paradoxical concept, suggesting that the brain codifies time by removing it, representing a dynamic situation as future critical interactions among elements encapsulated in a static map. According to the time compaction hypothesis, a dynamic situation is neurally represented as a compact internal representation (CIR, Fig. 1A). The CIR functions as a static map, spatially encoding all possible interactions between individual elements in a given environment, effectively removing time from the dynamic situation while retaining all the necessary information for navigation^20^. The existence of time compaction in humans has been recently demonstrated^21^.

**Figure 1:**
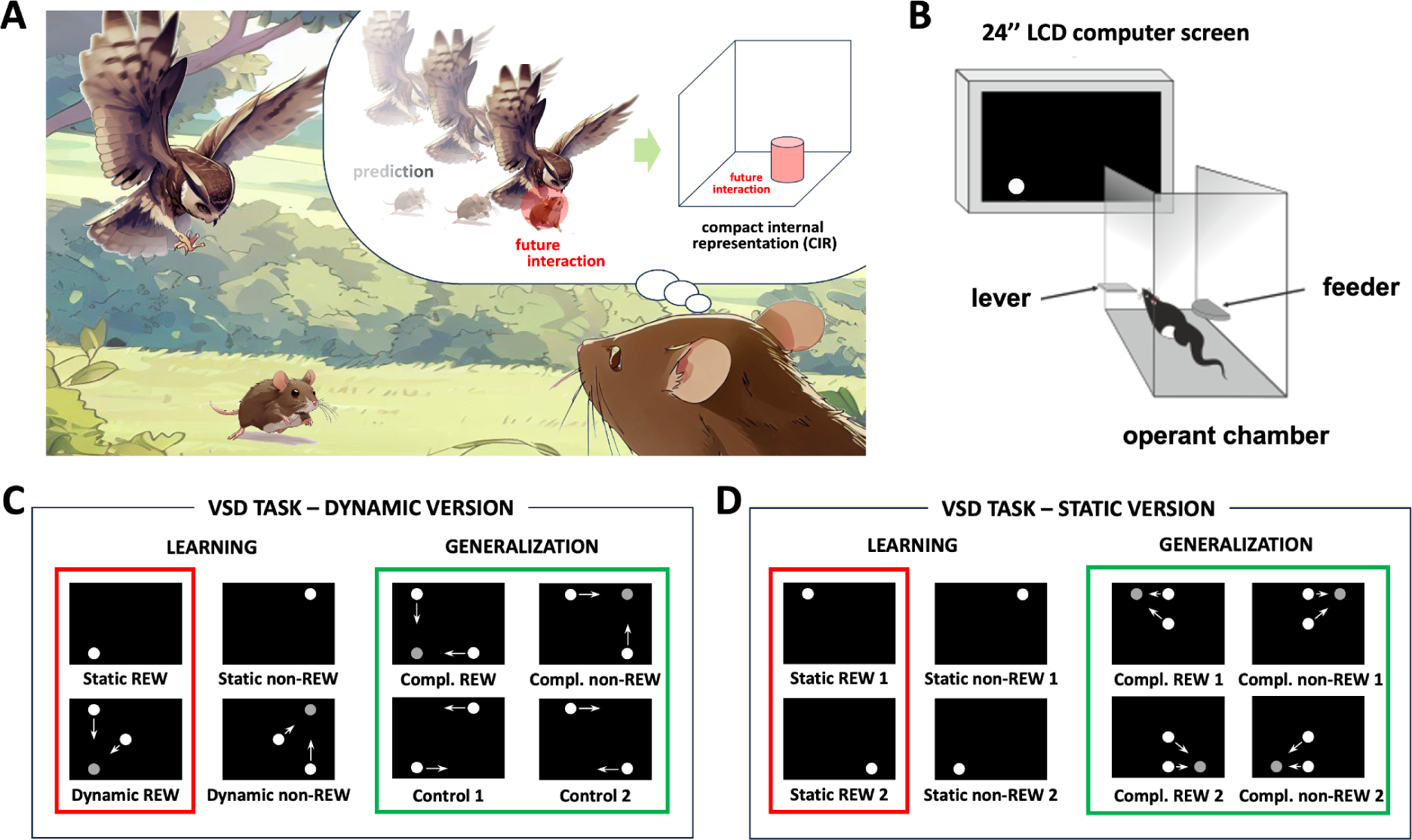
Experimental design. **(A)** Time compaction theory proposes that a dynamic scenario is internally represented by statically mapping the predicted future interactions. This static map, called compact internal representation (CIR), enables efficient learning and memorization of dynamic situations^20^, allowing real-time decision-making, a primary requirement for survival. **(B)** Visuospatial discrimination (VSD) task. We trained the rats to discriminate spatial stimuli on the distant computer screen. **(C, left)** VSD task - dynamic version. We presented the rats with two complementary rewarded stimuli – one static and one dynamic (Static REW and Dynamic REW, in red rectangle) and two complementary non-rewarded stimuli (Static non-REW and Dynamic non-REW). The static stimuli were stable white circles, and the complementary dynamic stimuli consisted of two circles moving toward the static circle location (grey circle – predicted collision point; not displayed) but disappearing half of the way. **(C, right)** During the generalization test related to the dynamic version of the VSD task, we presented the rat with four novel stimuli pseudorandomly displayed between the presentations of the four familiar training stimuli (shown in panel C, left). One novel stimulus shared the location of the predicted collision with the Dynamic REW stimulus (Compl. REW), another with the Dynamic non-REW stimulus (Compl. non-REW), and two more stimuli had no predicted collision point (Control 1 and Control 2). **(D, left)** VSD task - dynamic version. We used two rewarded static stimuli (Static REW1 and Static REW2, in red rectangle) and two non-rewarded static stimuli (Static non-REW1 and Static non-REW2) in this version of the VSD task. **(D, right)** During the generalization test related to the static version of the VSD task, we again presented the rat with four novel stimuli pseudorandomly displayed between the presentations of the four familiar training stimuli (shown in panel D, left). The novel dynamic stimuli had the location of the predicted collision identical to the static positions of familiar training stimuli: Compl. REW1 to Static REW1, Compl. REW2 to Static REW2, Compl. non-REW1 to Static non-REW1, Compl. non-REW2 to Static non- REW2.

We designed our study to understand the mechanisms of the internal representation of our spatiotemporal world, specifically the processing of visuospatial scenes. We 1) compared the discrimination of static and complementary dynamic visuospatial stimuli and 2) tested the time-compaction hypothesis in rats. Our results show that rats can internally represent dynamic visuospatial scenes as static maps capturing future interactions between objects in the environment, discriminating dynamic visuospatial stimuli faster and more precisely than static ones.

## METHODS

### Animals

The experimental subjects were outbred male Long-Evans rats (N=8 for the dynamic version of the task, N=6 for the static version) three months old at the beginning of the experiment. The rats were obtained from the Charles River Laboratories breeding colony (Calco, Italy; for the dynamic version of the task) or the Animal Facility of the Institute of Physiology CAS (Prague, Czech Republic; for the static version of the task) and were housed in pairs. The animal room was kept at a constant temperature of 21 °C and a 12-hour light/dark cycle. All handling and experiments were performed during the light phase of the day. Rats had free access to water but were limited to food to be maintained at 90% of their free-feeding weight. After transportation from the breeding colony, the rats were left to acclimatize for ten days and then handled for 5 mins every day for five days before the beginning of the behavioral training. Animal welfare complies with current legislation (the Animal Protection Code of the Czech Republic and EU Directive 2010/63/EC).

### Behavioral apparatus

The apparatus consisted of an operant chamber with a lever and a feeder, a 24’’ LCD monitor (1920×1080 pixels, 16:9 aspect ratio) placed 37 cm in front of the chamber, and a computer (Fig. 1B). The computer displayed the stimuli on the screen, registered lever presses, and activated the feeder. In the case of a rewarded lever press, a 45 mg chocolate-flavored pellet (Bio-Serv, USA) was delivered from the feeder to the operant chamber. The stimuli were displayed by software written in C++. During the experiments, rats were trained and tested in two identical apparatuses. Each rat was randomly assigned to a particular apparatus and always trained and tested in the same apparatus.

### Visuospatial discrimination (VSD) task - dynamic version

In the dynamic version of the VSD task, we trained the food-deprived rats (N=8) to discriminate by lever-pressing a particular static rewarded position of a circle displayed on a distant computer screen and a complementary dynamic rewarded stimulus consisting of two circles that approached but did not reach the same static position (VSD task - dynamic version, Fig. 1C). We progressively used five training configurations with increased demands on the rats’ performance. We gradually shortened the presentation of the stimuli from 90 s to 15 s and changed the reinforcement schedule from continuous to variable ratio 3. Individual visual stimuli lasted 15 s and were separated by 3 s blank screen periods in the final (fifth) configuration of the behavioral training. The complete behavioral protocol is described in Supplemental Methods and Supplemental Table 1.

We used two types of visual stimuli in the dynamic version of the VSD task: static and dynamic. The static stimuli were a white circle (d = 5.85 cm) in the screen’s bottom left or top right. The two complementary dynamic stimuli consisted of two such circles moving in the direction of the circles in the static stimuli, implying future collision at those positions, but with both moving circles disappearing in the middle of their trajectories. The starting position of the first moving circle was the top left or bottom right part of the screen, respectively, and the second moving circle’s starting position was the screen’s center (Fig. 1C, left). The duration of a single dynamic circle approach was 1 s. Therefore, the complete dynamic stimulus in the final configuration of the behavioral training, lasting 15 s, consisted of 15 repetitions of a particular approach. The laterally positioned circle was moving at 12.45 cm/s, and the circle that started in the center of the screen had a speed of 8.8 cm/s.

Of these four stimuli - two static and two dynamic - one static and the complementary dynamic were rewarded (Fig. 1C, left, marked in a red rectangle; Static REW, Dynamic REW), and the other static and its complementary dynamic version were non-rewarded (Fig. 1C, left; Static non-REW, Dynamic non-REW). One-half of the rats (N=4) were trained with rewarded and non-rewarded stimulus combinations shown in Fig. 1C. The other half of rats (N=4) were trained with the swapped combination of stimuli, i.e., the rewarded stimulus combination in the first half of the rats was the non-rewarded combination in the second half of the rats and vice versa. We used this design to exclude potential bias for a particular stimulus. For our analyses, we merged data from both halves of the rats as we did not observe any prominent sub-group-related differences in the discrimination performance of the trained rats.

### VSD task - dynamic version: Generalization test

Once the rats mastered the dynamic version of the VSD task, we conducted a generalization test. We introduced four novel dynamic spatial stimuli to examine the rats’ ability to apply their acquired knowledge of predicted collision points, whether rewarded or non-rewarded, to discern unfamiliar dynamic stimuli that shared the collision position with the familiar ones. We displayed the four novel stimuli pseudorandomly between the presentations of the four familiar training stimuli during the generalization test.

Among the four novel non-rewarded dynamic stimuli comprising two moving circles, two stimuli shared the collision position with the previously discriminated dynamic stimuli. Specifically, one stimulus aligned with the Dynamic REW stimulus (Compl. REW), while the other aligned with the Dynamic non-REW stimulus (Compl. non-REW). The remaining two dynamic stimuli, without the predicted collision point, were included as control stimuli to assess novelty preference (Control 1, Control 2) (Fig. 1C, right).

The hypothesis was that if the animals discriminated the dynamic spatial stimuli in the training phase based on the critical static information – the position of the predicted collision, they would prefer the novel dynamic stimulus that shares the position of the predicted collision with the previously discriminated Dynamic REW stimulus over the novel dynamic stimulus that shares the position of the predicted collision with the previously discriminated Dynamic non-REW stimulus, i.e., Compl. REW stimulus over Compl. non-REW stimulus. We performed the generalization test two times. Five standard training sessions separated both tests to eliminate the effect of the first generalization test on the rats’ performance in the second test.

### VSD task - static version

In the static version of the VSD task, we trained a different group of food-deprived rats (N=6) to discriminate two particular static rewarded positions of the white circle (Fig. 1D, left, marked in a red rectangle; Static REW1, Static REW2) and two static non-rewarded positions of the same circle (Fig. 1D, left; Static non-REW1, Static non-REW2). All aspects of the behavioral training and experimental design were the same as for the dynamic version of the VSD task described above. The only difference was the set of used stimuli. One-half of the rats (N=3) were again trained with rewarded and non-rewarded stimulus combinations shown in Fig. 1D, and the other half (N=3) were trained with the swapped combination of stimuli.

### VSD task - static version: Generalization test

After the rats mastered the VSD task’s static version, we performed a generalization test. The design of this generalization test was identical to the one described for the dynamic version of the task except for the set of presented stimuli. In this case, the novel dynamic stimuli placed the anticipated collision point exactly where the static positions of the familiar training stimuli were located: Compl. REW1 mirrored Static REW1, Compl. REW2 mirrored Static REW2, Compl. non-REW1 mirrored Static non-REW1, and Compl. non-REW2 mirrored Static non-REW2 (Fig. 1D, right). Therefore, according to the hypothesis, the rats were supposed to prefer Compl. REW stimuli over Compl. non-REW stimuli if they could represent the dynamic scene by the predicted collision point. We again performed the generalization test two times, and five standard training sessions separated both tests.

### Data analysis

For the dynamic version of the VSD task, we analyzed three aspects of rats’ behavior: learning dynamics, asymptotic performance, and stimulus generalization. We studied learning dynamics during the training sessions of the first training configuration and analyzed asymptotic performance in the last session of the final training configuration. At the same time, data from two generalization tests were used to analyze the possible transfer of the acquired knowledge to discriminate a set of novel dynamic stimuli.

Three main behavioral variables were analyzed to study rats’ performance during the presentation of individual stimuli: the mean probability of pressing the lever (mean number of lever presses per second), the probability distribution of lever-pressing during presentation periods (number of lever presses per ms within 200 ms bins over the time course of each presentation period), and the reaction time (time from the stimulus onset to the first lever press). Due to the lack of independence between data from the same animal, all models include a random intercept, taking the rat id as a grouping variable. Lever pressing probability was analyzed utilizing a generalized linear mixed model (GLMM) with binomial distribution and logit link function (i.e., a logistic regression model). In contrast, a Cox Proportional Hazards model studied the reaction time. In all cases, we included stimulus type as a fixed effect factor. For the probability distribution analysis, the time from the stimulus onset (binned into 200 ms periods), its square, and its interactions with the stimulus type were also included as fixed effects factors. Analysis involving learning dynamics included the training duration, i.e., the number of sessions within each training configuration normalized for each rat by the maximum number of sessions that a rat underwent to reach the learning criteria of a particular configuration, as a fixed effect factor (both alone and in interaction with the stimulus type) and a random slope.

For the static version of the VSD task, we analyzed only the two generalization sessions in the same way as described above for the dynamic version of the VSD task. As rats were trained with only static stimuli, the stimulus type factor for models had two levels (Static REW/Static non-REW for the familiar stimuli and Compl. REW/Compl. non-REW for the novel stimuli, respectively) instead of four levels as in the generalization session of the dynamic version of the VSD task.

A tentative full model including all factors and interactions was carried out in all regression models. Then, we selected variables by a stepwise backward procedure using *p*-values and AIC as criteria to exclude them until we reached a model with all significant factors. To assess differences between levels of significant categorical factors (stimuli types), type II Wald Chi-Square and Tukey posthoc tests with FDR correction were done. The level of significance was set to 0.05 in all cases. Data are shown as mean ± standard error of the mean (SEM). For a more detailed description of the studied variables, justification of the models used, and interpretation of the model’s coefficients, see Supplemental Methods.

We carried out all analyses in R (version 4.1.3^22^) under RStudio (version 2022.02.3^23^). We used R packages lme4^24^, emmeans^25^, survival^26^, coxme^27^, and ggplot2^28^.

## RESULTS

### VSD TASK - DYNAMIC VERSION

In the dynamic version of the VSD task, we analyzed both the learning phase and the subsequent generalization phase. The data from the learning phase demonstrate the better discrimination of dynamic versus static visuospatial stimuli. On the other hand, the data from the generalization phase suggest that the rats can represent dynamic environments by static maps containing future interactions of moving objects.

### Learning phase

In nature, threads and opportunities for survival are essentially dynamic. Thus, animals must primarily deal with dynamic scenarios requiring critical real-time decisions. Therefore, we hypothesize that dynamic stimuli are prioritized over static stimuli in central cognitive processes such as learning. To test this hypothesis, we conducted a visuospatial discrimination task in which rats were trained to discriminate static and complementary dynamic visuospatial stimuli.

### First learning configuration: faster learning and shorter reaction times of dynamic versus static stimulus discrimination

We assessed rats’ learning performance by the probability of pressing the lever during the presentation of particular stimuli. During the first training configuration, the probability of lever pressing increased with the training progression for rewarded stimuli while decreased for non-rewarded ones (p < 0.0001 for all four slopes). Although there were no differences between slopes of non-rewarded stimuli (p = 0.31), lever pressing probability increased faster during Dynamic-REW than during Static-REW stimulus (p = 0.005), so at the end of the first training configuration, this probability was 30% higher for dynamic than for static rewarded stimulus (odds ratio (OR) = 1.32, Fig. 2A). On the other hand, the hazard ratio analysis showed that the reaction time decreased with training progress for rewarded stimuli while increased for non-rewarded stimuli (p < 0.0001 for all comparisons of any rewarded stimulus with any non-rewarded stimulus) (Fig. 2B). While the probability of pressing the lever for the first time at any time from the onset of the stimulus is 19% higher in Dynamic-REW compared with Static-REW (p = 0.01, HR = 1.19), there were no differences between non-rewarded stimuli (p = 0.06, HR =1.09). These findings suggest faster learning of dynamic visual stimuli than similar static stimuli.

**Figure 2:**
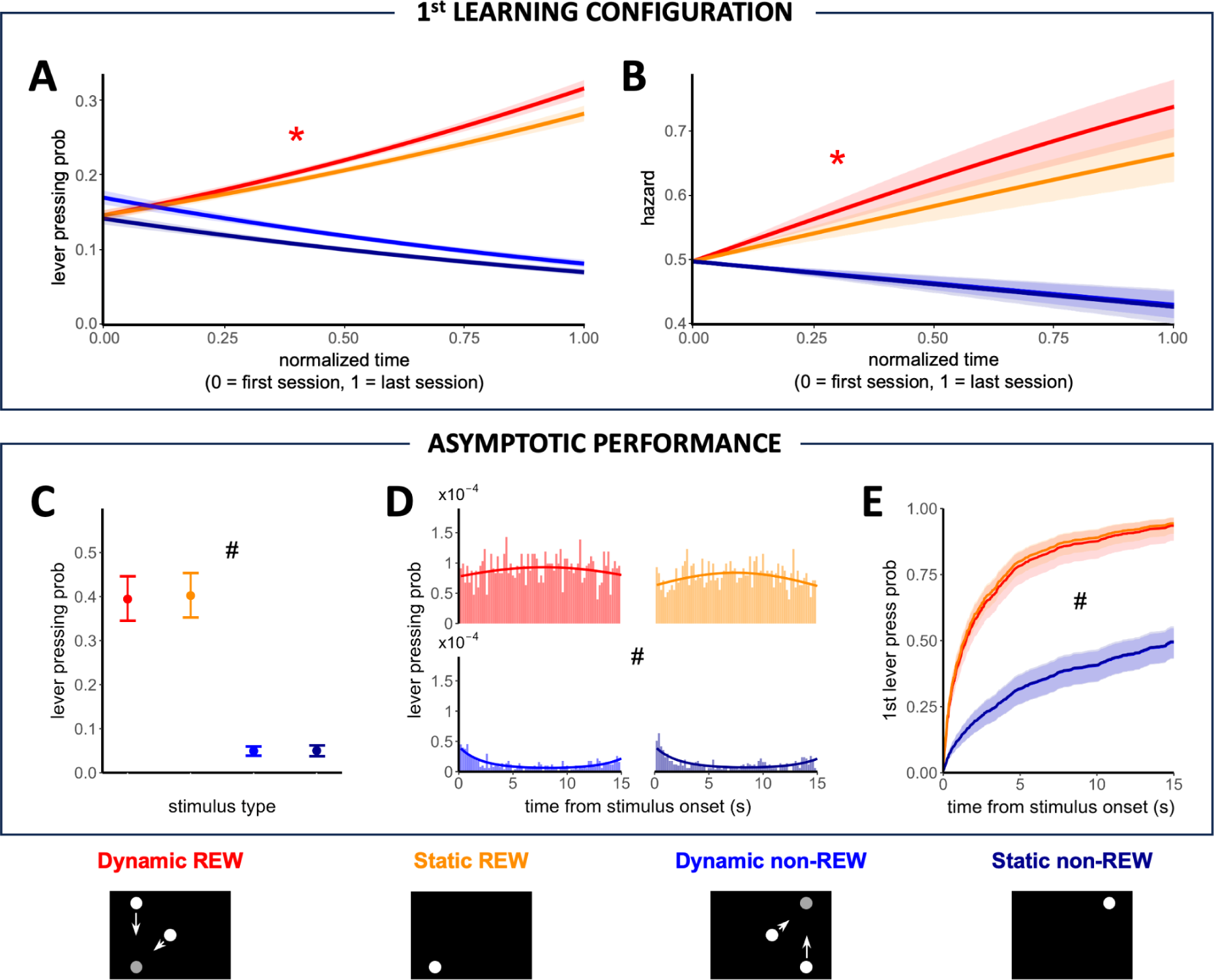
Discrimination of dynamic and static visuospatial stimuli. *Faster learning and shorter reaction times of dynamic versus static stimulus discrimination.* **(A)** Lever-pressing probability for individual stimuli during the first training configuration. **(B)** Hazard ratios of the first lever press after the stimulus onset for individual stimuli during the first training configuration. *Equal discimination of dynamic and static stimulus during asymptotic performance.* **(C)** Mean probability of pressing the lever within 1-s bins of the presentations of individual stimuli during the last session of the final training configuration. **(D)** The probability distribution within 1-ms bins throughout individual stimuli duration (histograms binned at 200 ms). **(E)** The probability of the first lever press after the stimulus onset for individual stimuli during the last session of the final training configuration. The red asterisk marks significant differences (p < 0.01) between the Dynamic REW stimulus and all other stimuli, and the black hashtag marks significant differences (p <0.0001) between rewarded and non-rewarded stimuli. Further statistical differences are described in the text. Data in panels C and E are shown as means ± SEM.

As individual rats underwent different numbers of sessions during discrete training configurations (Suppl. Tab. 2), we normalized the learning time for individual configurations to a scale from 0 (first session in the particular training configuration) to 1 (last session in the specific training configuration). Learning curves for all four stimuli were prominent only during the first configuration of the task (Fig. 2A). During the subsequent configurations, in which we gradually shortened the presentation time of individual stimuli and increased the schedule of reinforcement, learning curves were not significant or selective for a particular stimulus (Suppl. Fig. 1). The primary purpose of these subsequent configurations was to increase the demands on attention during the task and prevent the rats from using the feedback of the apparatus (reward delivery) to solve the task. Out of the eight rats trained in the dynamic version of the VSD task, seven reached a stable performance of > 60% in the final configuration of the task. On average, the rats took 46 (± 2.4 SEM) sessions to be fully trained in the task (see Supplemental Table 2 for more details).

### Asymptotic performance: equal discrimination performance for dynamic and static stimuli

In the last session of the final training configuration, when the rats had already reached the asymptotic performance in the VSD task (see Suppl. Figs. 1-3 for more details about the rats’ performance during training configurations 2-5), the lever-pressing probability during the rewarded stimuli was higher than during the non-rewarded stimuli (p < 0.0001 in all cases). However, as expected, there was no difference between the two rewarded (Dynamic REW vs. Static REW, p = 0.96) or non-rewarded stimuli (Dynamic non-REW vs. Static non-REW, p = 1) (Fig. 2C-2D).

**Figure 3:**
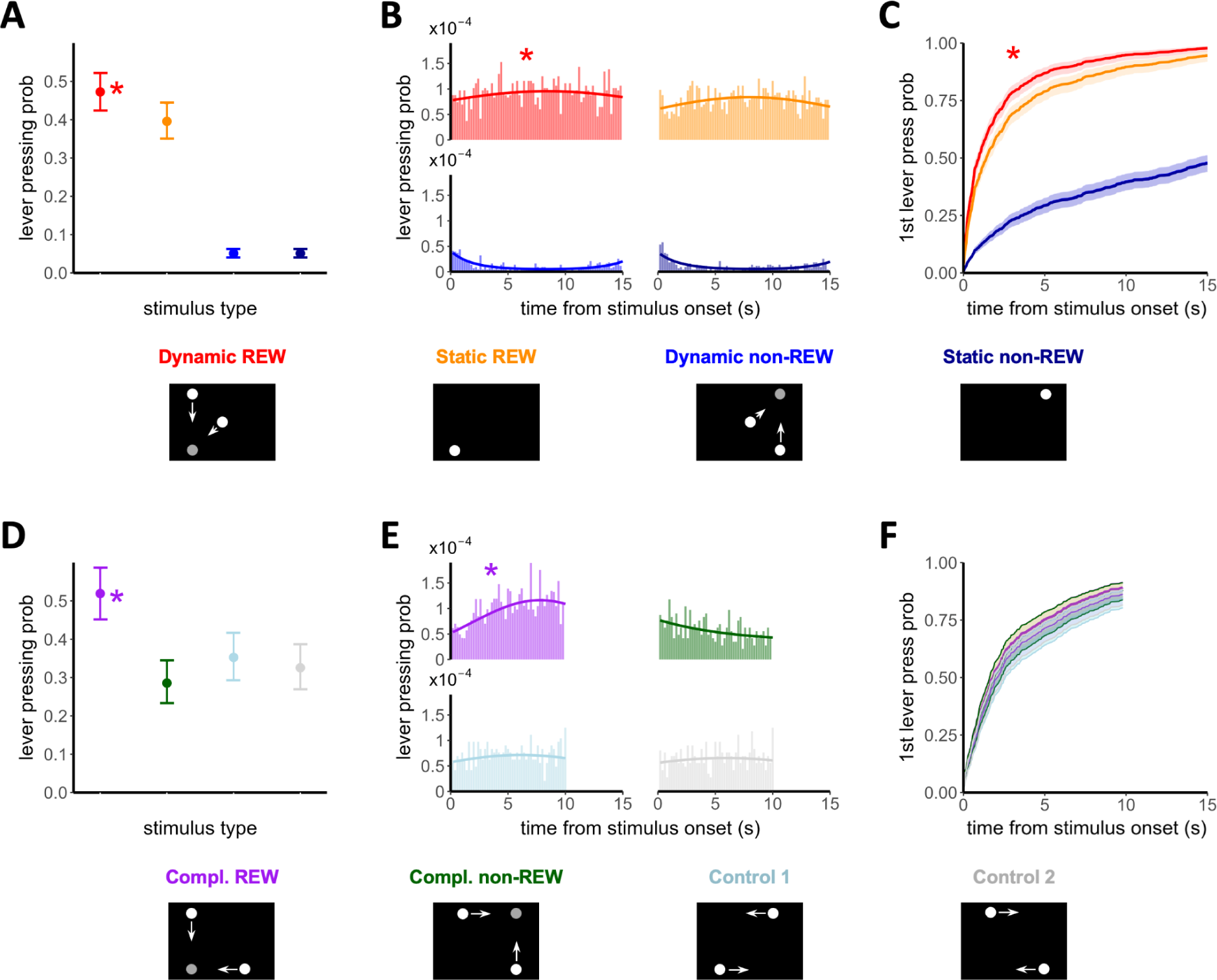
First evidence of the internal representation of future interactions in rats. The rats reinstated their preference for the rewarded dynamic stimulus over the static one when novel non-rewarded dynamic stimuli were introduced. (A) Mean probability of pressing the lever within 1 s bins of the presentations of individual familiar stimuli during the generalization test. (B) The probability distribution within 1-ms bins throughout stimuli duration (histograms binned at 200 ms). (C) The probability of the first lever press after the stimulus onset for individual familiar stimuli. The red asterisk marks significant differences (p < 0.0001) between the Dynamic REW stimulus and all other novel stimuli; further statistical differences are described in the text. *The rats preferred the novel dynamic stimulus in which two circles approached the collision point in the familiar dynamic stimulus and the familiar static reward position.* (D) Main probability of pressing the lever within 1 s bins of the presentations of individual novel stimuli during the generalization phase. (E) probability distribution within 1 ms bins throughout stimuli duration (histogram binned at 200 ms). (F) The probability of the first lever press after the stimulus onset for individual novel stimuli during the generalization tests. The purple asterisk marks significant differences (p < 0.0001) between the Compl. REW stimulus and all other novel stimuli; further statistical differences are described in the text. Data in panels A, C, D, and F are shown as means ± SEM.

Visual stimulus type affected the time to the first lever press after the stimulus onset (p < 0.0001; Fig. 2E). Tukey posthoc tests showed a shorter time to the first operant response for the rewarded stimuli than for the non-rewarded stimuli (all p < 0.0001). Again, there was no difference between the two rewarded (p = 0.83) or non-rewarded stimuli (p = 1). The median time to the first lever press after the stimulus onset was 1760 ms and 2040 ms for the Dynamic REW and Static REW stimulus, respectively. The median time was longer than the stimulus duration for both non-rewarded stimuli. This lack of differences between Dynamic REW and Static REW suggests that after extensive training, the rats can discriminate similar dynamic and static visuospatial stimuli equally.

### Generalization phase

The existence of a compact internal representation (CIR) of predicted interactions can be tested by the visuospatial discrimination of appropriate stimuli that share critical spatial information. The generalization phase of the dynamic version of the VSD task utilizes novel dynamic stimuli sharing the point of future collision with the familiar stimuli from the learning phase. According to the time compaction hypothesis, such complementary familiar and novel stimuli should be represented by similar CIRs. Thus, we expected that the rats previously trained to discriminate a particular stimulus would prefer a novel stimulus when both share the locations of future interactions.

### Reinstated preference for rewarded dynamic over static stimulus when novel non-rewarded dynamic stimuli were introduced

During the generalization phase, the analysis concerning learned stimuli revealed higher lever-pressing probability during the Dynamic REW stimulus than during all other familiar stimuli (all p < 0.0001) and during the Static REW stimulus than during both non-rewarded stimuli (both p < 0.0001). There was no difference between the two familiar non-rewarded stimuli (p = 0.773) (Fig. 3A-3B). The probability of lever-pressing during the presentation of the rewarded dynamic stimulus was 18% higher than while the rewarded static stimulus was displayed (odds ratio (OR) = 1.19).

Visual stimulus type again affected the time to the first lever press after the stimulus onset (p < 0.0001; Fig. 3C). Contrary to the last training session, Tukey posthoc tests showed a shorter time to the first operant response for the Dynamic REW stimulus than for the three other stimuli (all p < 0.0001). As in the last training session, there was a shorter time for the first lever press for the Static REW stimulus than for both non-rewarded stimuli (p < 0.0001). There were no differences between the two non-rewarded stimuli (p = 0.999). The hazard ratio (HR) for the Dynamic REW stimulus was 1.3 times higher than for the Static REW, meaning that at any time after the onset of the stimulus, the probability that the lever has already been pressed was 30% higher for the dynamic rewarded stimulus than for the static ones. In other words, during the Dynamic REW stimulus, the rats pressed the lever for the first time earlier than during the Static REW stimulus (median time of 1360 ms and 2000 ms for the Dynamic REW and Static REW stimulus, respectively; the median time was longer than the stimulus duration for both non-rewarded stimuli). Note that both generalization sessions (see Methods) were combined for the analysis since the regression models did not reveal any influence of this factor.

### Evidence of the CIR: preference for the novel dynamic stimulus that shares the spatial information with rewarded familiar stimuli

To precisely evaluate the differences among individual novel stimuli, we analyzed the non-familiar stimuli used in the generalization tests separately from the familiar stimuli. This approach aimed to eliminate any potential influence of novelty on the lever-pressing activity.

The mean probability of pressing the lever was significantly higher during the Compl. REW stimulus, i.e., the dynamic stimulus where the point of future collision coincides with that in the familiar dynamic stimulus (Dynamic REW) and with the position of the circle in the familiar static stimulus (Static REW), than during other novel stimuli (all p < 0.0001, Fig. 3D). This probability for the Compl. REW stimulus is about 79% higher than during the Compl. non-REW stimulus (odds ratio (OR) = 2.7) and 49% and 58% higher than during the Control 1 and Control 2 stimuli, respectively (odds ratio (OR) = 1.98, odds ratio (OR) = 2.24, respectively). The lever pressing probability was significantly lower during the Compl. non-REW stimulus than during the Control 1 stimulus (20% lower, p < 0.0001) and lower, but not significantly, than during the Control 2 stimulus (10% lower, p = 0.06). Both control stimuli showed similar probabilities (p = 0.11).

Since the interaction between the stimulus type and time was significant (p < 0.0001), the probability of pressing the lever throughout the presentation of individual stimuli was different between the stimuli (Fig. 3E). The rats preferred the novel dynamic stimulus that shares the position of the predicted collision with the previously discriminated Dynamic REW stimulus, i.e., the Compl. REW stimulus, over the other three novel stimuli. None of the control stimuli showed a significant dependence on time, neither the linear nor the quadratic component, while the Compl. non-REW stimulus (the stimulus that shares the place of the predicted collision with the previously discriminated Dynamic non-REW stimulus) depended only on the linear component of time with a negative slope. Both linear and quadratic components of time had a significant effect (p < 0.0001 in both cases) on the probability of pressing the lever during the presentation of the Compl. REW stimulus.

In contrast to the familiar stimuli, there was no difference between the time at which the rats pressed the lever for the first time after each stimulus onset among the novel stimuli in the generalization tests (p = 0.2347; Fig. 3F). The lever was pressed for the first time after the stimulus onset at similar times for all four stimuli (median times of 1920 ms, 1520 ms, 1860 ms, and 1960 ms for the Compl. REW, Compl. non-REW, Control 1, and Control 2 stimulus, respectively).

### VSD TASK - STATIC VERSION

In the static version of the VSD task, we analyzed solely the generalization phase in previously well-trained rats. In this case, we displayed dynamic stimuli to the rats for the first time, and those stimuli had no visual overlap with the familiar stimuli as they did during the generalization tests of the dynamic version of the VSD task. These results confirm the rats’ ability to represent the future interaction of two moving objects by their static intersection.

### Generalization phase

The Compl. REW and Dynamic REW, and the Compl. non-REW and Dynamic non-REW stimuli, respectively, share not only the predicted collision point but also one of the moving circles in the dynamic version of the VSD task (Fig. 1C). Therefore, to confirm the previous indications of the CIR existence, we conducted a second generalization experiment where the learned familiar stimuli are purely static and the only shared information with the novel moving generalization stimuli involves the location of their future interactions.

### Evidence of the CIR: preference for the novel dynamic stimuli in which two circles approached the familiar static reward positions

The static version of the task served the sole purpose of conducting a different version of the generalization test. In this test, we introduced dynamic stimuli to the rats for the first time, enabling us to assess the time compaction mechanism without the risk of partial visuospatial information transfer (for more details, see the Discussion section).

As stated in Methods, both types of stimuli (REW and non-REW) were counterbalanced between the screen’s top/bottom and left/right sides. Regression models did not show any effect of these factors on the dynamics of lever pressing behavior. Thus, to avoid any bias, they were combined so that the factor stimulus type only has two levels, REW and non-REW. As in the case of the dynamic version of the VSD task, both generalization sessions were combined for analysis. During the generalization phase, the analysis revealed a higher lever-pressing probability during the Static REW stimuli than during the Static non-REW stimuli (p < 0.0001) (Fig. 4A-4B). The probability of lever-pressing during the presentation of the rewarded stimuli was 420% higher than while the non-rewarded stimuli were displayed (odds ratio (OR) = 5.22).

**Figure 4:**
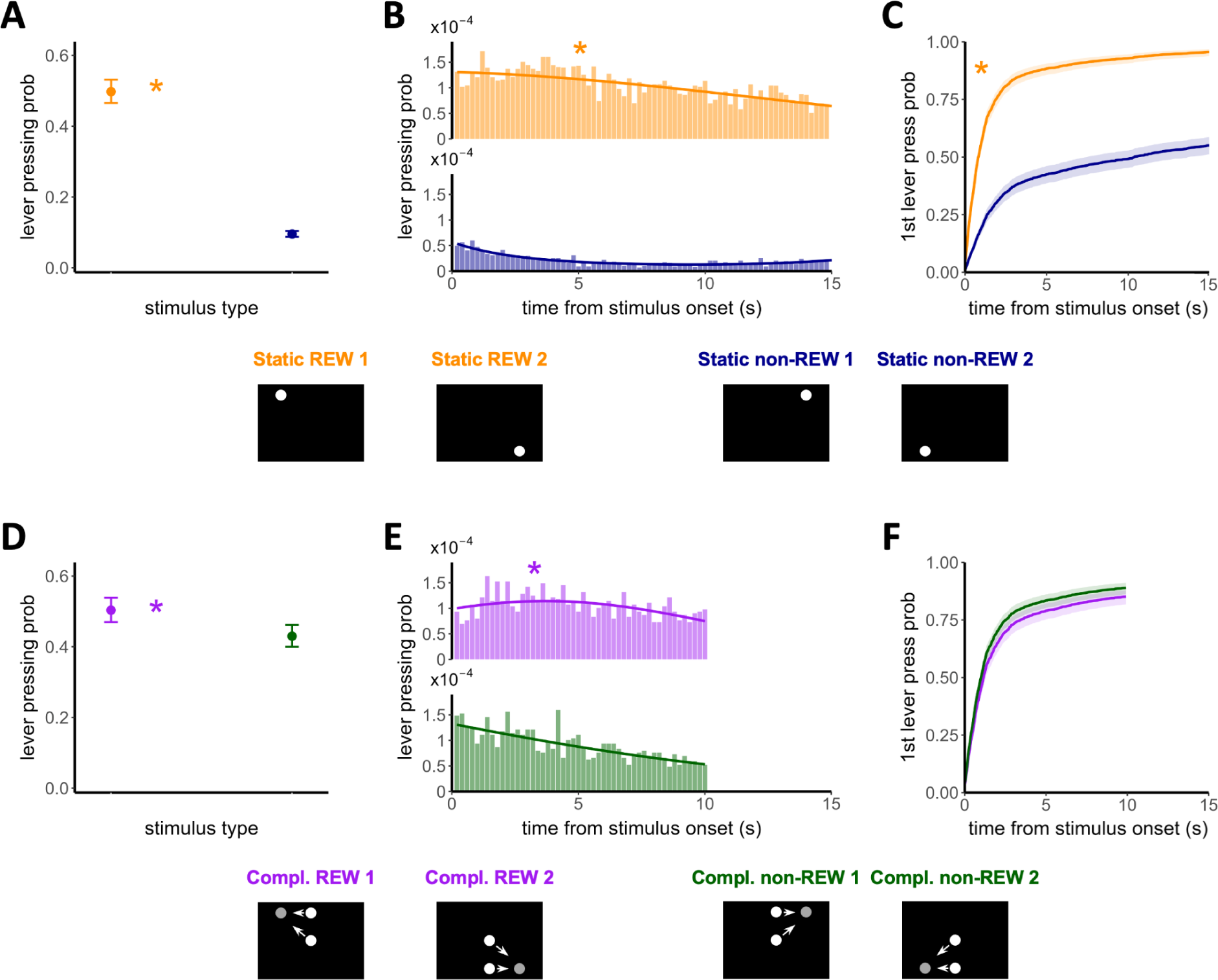
Confirmation of the internal representation of future interactions in rats. *The rats discriminated between rewarded and non-rewarded familiar static object positions.* **(A)** Mean probability of pressing the lever within 1 s bins of the presentations of the two types of familiar stimuli during the generalization test. **(B)** The probability distribution within 1-ms bins throughout stimuli duration (histograms binned at 200 ms). **(C)** The probability of the first lever press after the stimulus onset for both types of familiar stimuli. The orange asterisk marks significant differences (p < 0.0001). *The rats preferred the novel dynamic stimuli in which two circles approached the familiar static reward positions*. **(D)** Main probability of pressing the lever within 1 s bins of the presentations of the two types of novel stimuli during the generalization phase. **(E)** probability distribution within 1 ms bins throughout stimuli duration (histogram binned at 200 ms). **(F)** The probability of the first lever press after the stimulus onset for both types of novel stimuli during the generalization tests. The purple asterisk marks significant differences (p < 0.0001). Data in panels A, C, D, and F are shown as means ± SEM.

Visual stimulus type again affected the time to the first lever press after the stimulus onset (Fig. 4C), so the Cox Regression model showed a shorter time to the first operant response for the Static REW stimuli than for the Static non-REW stimuli (p < 0.0001). The hazard ratio (HR) for the Static REW stimuli was 2.82 times higher than for the Static non-REW stimuli, meaning that at any time after the onset of the stimulus, the probability that the lever has already been pressed was 180% higher for the Static REW stimuli than for the Static non-REW stimuli (median time of 1000 ms for the Static REW stimuli and 11640 ms for the Static non-REW stimuli, respectively).

The mean probability of pressing the lever was higher during the Compl. REW stimuli than during Compl. non-REW stimuli (p < 0.0001, Fig. 4D). Since the interaction between the stimulus type and time was significant (p < 0.0001), the probability of pressing the lever was also different throughout the presentation of stimuli (Fig. 4E). The rats preferred the novel dynamic stimuli that share the position of the predicted collision with the previously discriminated Static REW stimuli. The probability of lever-pressing during the presentation of the Compl. REW stimuli were 17% higher than while the Compl. non-REW stimuli were displayed (odds ratio (OR) = 1.17).

There was no difference in the time at which the lever was pressed for the first time after the stimulus onset between the novel stimuli in the generalization tests (p = 0.30; Fig. 4F). The lever was pressed for the first time after the stimulus onset at similar times for both types of stimuli (median times of 1280 ms and 800 ms for the Compl. REW and Compl. non-REW stimuli, respectively).

## DISCUSSION

Nature is essentially dynamic, and survival requires efficient visual information processing to make real-time critical decisions and cope with threats and demands. In this work, we present data obtained from behavioral experiments in rats supporting the preferential processing of dynamic stimuli and the involvement of the previously introduced time compaction mechanism in representing dynamic environments.

First, we showed that rats discriminate dynamic visual objects faster and more precisely than static visual objects. Second, our results suggest that rats can effectively use the time compaction mechanism to internally represent dynamic spatial situations as static maps capturing predicted object interactions. This work supports the encoding of future interactions as an efficient mechanism of spatiotemporal cognition, suggesting it as an evolutionary invariant after similar phenomena are reported in humans^21^ and bats^29^. Moreover, spatiotemporal processing is essential for episodic memory^30^, and its impairment is a precursor to cognitive decline and has consequences for various neuropsychiatric illnesses, including dementia and schizophrenia^31-34^. Therefore, the obtained information could open new venues for creating innovative therapeutic strategies to combat cognitive deterioration. In addition, developing transparent and interpretable artificial navigation systems and robots with animal-like navigation abilities can also be aided by understanding the mechanics underlying spatiotemporal cognition in dynamic environments^35,36^.

We designed unique behavioral experiments to test rats’ processing of static and dynamic objects outside their direct reach. The dynamic nature of objects and their inaccessibility are characteristics typical of natural conditions that were overlooked in previous studies. Furthermore, we presented the rats with static and dynamic stimuli that were complementary in their spatial aspects, allowing their direct comparison.

### Rats learn to discriminate dynamic visuospatial stimuli faster, with better accuracy and shorter reaction time than static visuospatial stimuli

Acting swiftly and precisely in complex dynamic surroundings is an essential ability. Natural ecosystems are constantly changing and responding especially to dynamic elements of the environment (e.g., an approaching predator, a falling boulder, etc.) is crucial. At the same time, static objects usually do not pose an immediate threat. Therefore, processing primarily dynamic events quickly and effectively would be beneficial from an ecological standpoint.

In the learning phase of the experiment, we found that the rats learned to discriminate the dynamic stimuli faster, more precisely, and with shorter reaction times than the static ones. Our results agree with previous human studies reporting that participants detect moving objects more accurately than their static versions^37^ and that drivers detect moving objects sooner than static objects^38^. In addition, greater attention to moving objects than static objects was observed in human infants^39^, adult human subjects^40,41^, and mice^42^.

Abrams and Christ ^40^ proposed that the onset of movement, not the movement itself, captures attention. In our task, the dynamic stimuli consisted of 1-s long repetitions of individual approaches of two moving circles. Thus, the movement of the objects was repeatedly initiated during the dynamic stimuli presentations. This could attract even more attention to the dynamic stimuli and affect the rats’ performance in our task. However, we observed faster responses to the dynamic stimulus after its onset compared to the static stimulus only for the pair of rewarded stimuli but not for the pair of non-rewarded stimuli (Fig. 2B). Nevertheless, given that the onset of movement is the critical period for identifying potential threats from moving objects, it may be essential to pay attention to it^40^. In another study, Smith and Abrams^43^ conducted a series of experiments that confirmed attentional capturing by motion onset in humans and demonstrated its validity in two distinct scenarios: the animated movement of objects on a computer screen and the natural motion of real objects.

On the other hand, the higher complexity of dynamic stimuli in comparison to static stimuli in our VSD task (two objects vs. one object) was unlikely to contribute to faster learning as it was previously shown that rats learn to discriminate concurrent complex visual scenes with multiple objects with the same performance as concurrent single visual objects^44^.

Altogether, these findings suggest that the ability to distinguish moving stimuli with superior performance, compared to static stimuli, could be a phenomenon shared among different species and an evolutionary advantage critical for survival in the ever-changing world.

### Rats can represent dynamic environments by predicting interactions of moving elements

The mechanisms of processing complex dynamic environments are unknown. In pigeons, neurons that encode the time^45,46^ or distance^47^ to the collision of objects approaching the animal were discovered. However, to effectively navigate natural environments, animals must not process only potential encounters with looming objects but also interactions of and with other moving objects in their surroundings.

The generalization tests of our VSD task showed that the rats could transfer previously acquired knowledge and generalize novel visuospatial stimuli. They preferred the novel dynamic stimuli in which the two circles approached the familiar rewarded static positions or the position of predicted collision of circles in the familiar rewarded dynamic stimulus.

In the generalization tests of the dynamic version of the VSD task, the design of the familiar and novel stimuli raises a possible explanation of whether the rats’ generalization performance could be based purely on the direct transference of partial visuospatial information: the familiar Dynamic REW stimulus and the novel Compl. REW stimulus share the circle moving vertically from top to bottom (Fig. 1C). This common feature could eventually trigger the lever-pressing behavior when the Compl.

REW stimulus was presented, explaining the reported preference for it among novel stimuli. The lever-pressing probability for this stimulus during the generalization tests is equal (data not shown) to the probability for the Dynamic REW stimulus (Figs. 3A, 3D). However, their performance significantly drops when rats discriminate novel visual scenes in which part of the familiar rewarded stimulus is relocated^48^, the stimulus is turned upside down^49^, or when discriminating novel visual shapes, which were, in fact, partially rearranged familiar rewarded stimuli^50^. The novel stimuli in our generalization tests of the dynamic version of the VSD task are also partially rearranged or relocated familiar stimuli. Therefore, based on the previous findings, we should expect lower lever-pressing probability during the presentation of the Compl. REW stimulus in comparison to the Dynamic REW stimulus in our task. However, this was not the case, suggesting that the information transference by the partial visual similarity between Dynamic REW and Compl. REW stimuli was not the only factor operating in the generalization tests. We propose that another mechanism, compatible with encoding future interactions hypothesized by the time compaction, is involved during the discrimination of novel dynamic stimuli in our VSD task.

This explanation was supported by the results of generalization tests of the static version of the VSD task, where the novel stimuli did not overlap the visual information with the familiar stimuli. Indeed, the rats were presented for the first time with a set of dynamic stimuli consisting of two moving circles with a predicted collision point. This location of the future collision was the only mutual information between the familiar static and novel dynamic stimuli. Still, the rats could generalize the novel dynamic stimuli and preferred the stimuli with the predicted collision point at the positions of familiar rewarded stimuli. These results suggest that this location of future interaction is crucial for discriminating and generalizing the dynamic visuospatial stimuli in our VSD task.

### Rats can use the time compaction mechanism to represent dynamic visuospatial environments

It was previously shown that rats can generalize static visual objects based on their spatial characteristics – specifically by estimating the distance between the familiar object and the novel object’s positions^14^ or the degree of geometric similarity between objects displayed at the same location^50^. A mechanism called time compaction has been introduced to decipher how time-changing environments could be encoded in the brain^19^. The inclusion of the time dimension in processing dynamic situations introduces a significant amount of redundant information, which has the potential to hinder the speed and accuracy of neural processing of individual events. This might be problematic, especially under critical life-threatening circumstances. Therefore, it would be beneficial for complex dynamic environments to be internally represented simply, enabling faster processing. Time compaction proposes that the brain embeds time into space rather than directly encoding it. This way, spatial dynamic situations are internally processed as compact internal representations (CIRs) – static maps containing the possible interactions between elements in a given environment. Since the CIRs encode future interactions between the environmental elements, they relate to the subject’s ability to predict the future positions of moving objects^21^. Following the time compaction hypothesis and the obtained results, we propose that rats could predict the future positions of moving objects in our task, imagine their potential collisions, and use the time compaction mechanism to generalize the novel stimuli in the generalization tests. Future subjects’ positions and movement directions are represented by distinct neural mechanisms nested in the hippocampus – a brain structure that plays a crucial role in spatial cognition. Several neuronal types have been discovered within the rodent hippocampus that give rise to a mental representation of the environment, known as a cognitive map^1^. These include the hippocampal place cells that encode the animal’s position in space^51^. Theta phase precession, a phenomenon that ties together the firing of place cells with the typical hippocampal theta rhythm, has been shown to predict movement direction^52^. Moreover, place cells have been found to fire in time-compressed sequences that predict one’s future trajectory during the periods of sharp wave ripples, the high-frequency oscillatory patterns presented in the mammalian hippocampus^53,54^. This way, future animals’ trajectories are pre-played in the brain within a few hundred milliseconds, while the real path lasts a few seconds. It suggests that the minimalization of the time dimension might be a general mechanism for predictive representation in the brain.

In addition, hippocampal cells in freely flying bats encode their future positions strongly related to intersections between trajectories^29^, as predicted by the time compaction hypothesis. In the Dotson and Yarstev study^29^, the neurons fired concerning nonlocal subjects’ positions in the environment. They represented the future locations of the bats, both during random foraging and goal-directed navigation. Moreover, most of these cells had their firing fields located at the intersections of the flying paths. These intersections correspond to the CIRs proposed by the time compaction mechanism. Their overrepresentation within the hippocampus suggests their high importance for navigation, and the results obtained in our study indicate a similar role for the predicted collision locations of moving objects for representing dynamic visual environments.

Our findings reveal that the rat’s brain can create abstract representations of inaccessible spaces where future interactions are spatially arranged. This is one of the central predictions of the time compaction hypothesis. Its experimental confirmation makes plausible the existence of the compact internal representation as a cognitive mechanism to deal with dynamic situations. In general, according to the time compaction hypothesis, when the subject is part of a dynamic situation (e.g., when it moves among other subjects) the CIR is a static map containing, besides the predicted interactions, the potential actions to deal with the situation (i.e., the pathways to be followed)^19^. In the same way, conventional cognitive maps mentally represent static environments by spatially mapping actual objects and trajectories to be navigated^53,55,56^. Thus, considering that the position of a static object will be the location of the future potential interactions of the agent with the object, the CIR would be a generalization of the cognitive map for time-changing scenarios^35^. Our results suggest a single general mechanism to represent dynamic and static situations mentally. This opens a new venue to the study of spatiotemporal cognition, as the time compaction hypothesis claims that the same neural populations responsible for processing static spaces would be involved in processing dynamic situations by spatially representing future interactions.

Recent theories propose that cognitive maps do not represent only explicit maps of space but can predict possible future states, e.g., possible future rewards or spatial positions in the environment^57^. The positional information about future location may be more critical for survival than the current position. Predictive cognitive maps could utilize successor representations^58^. Both rats and humans demonstrate analogous navigation choices and trajectory patterns, mirroring the behavior of reinforcement learning agents utilizing successor representation to adapt effectively to changing environments^59^. These predictive capabilities of the hippocampus and its central role in generating cognitive maps and processing static and dynamic situations, make this area the prime candidate for characterizing the cell populations responsible for the CIR.

In summary, we conclude that rats discriminate dynamic visuospatial stimuli faster and more precisely than static ones. Most importantly, our results suggest that rats can represent dynamic environments by creating static maps that capture significant expected interactions between objects. We emphasize the necessity of time-compacted spatial prediction in navigating complex dynamic environments requiring quick decision-making.

## ACKNOWLEDGMENTS

This research was supported by the project National Institute for Neurology Research (Programme EXCELES, ID Project No. LX22NPO5107)—Funded by the European Union–Next Generation EU, the Ministry of Education, Youth and Sports of the Czech Republic (MSMT) programme OP VVV project FGÚ MSCA Mobilita IV CZ.02.2.69/0.0/0.0/20_079/0017164, and Czech Science Foundation grant GACR 21-16667K. The authors from Spain were supported by the Ministry of Science and Innovation (Spain), Grant PID2022-138659NB-I00.

## AUTHOR CONTRIBUTIONS

T.D., A.S-J., J.A.V-A. and D.L. designed the experiments; T.D. and D.L. conducted the experiments and collected the data; A.S-J., P.M-C., J.A.V-A. and D.L. analysed and interpreted the data; V.L. created new software used in the work; T.D., A.S-J., J.A.V-A. and D.L. drafted the manuscript; all authors revised the manuscript; D.L., A. S. and J.A.V-A. provided the leadership and acquired funding.

## DATA AVAILABILITY STATEMENT

The datasets generated during and/or analysed during the current study are available from the corresponding author on reasonable request.

