## Supplemental information for "Internal representation of future interactions in rats"

### SUPPLEMENTAL METHODS

#### Visuospatial discrimination (VSD) task

Before starting the behavioral training, the food-deprived rats were pre-trained to press the lever in the operant chamber to obtain a food pellet. Then, we trained the rats once a day in the VSD task, five days a week, until they reached the performance criteria in the final training configuration of the task.

In the initial training configuration, individual stimuli were displayed for 90 s and were separated by 5 s blank screen pauses. As the single dynamic circle approach lasted 1 s, the complete dynamic stimulus consisted of 90 repetitions of the particular approach. We trained the rats under the continuous reinforcement schedule. Both rewarded stimuli were displayed three times, and both non-rewarded stimuli six times during the initial training sessions, which lasted 28 min 30 s. The sequence of the stimuli was pseudorandom to prevent the rats from using alternative strategies to solve the task. In addition, the same stimulus was never displayed twice in a row, and neither were two rewarded stimuli. As the non-rewarded stimuli were presented twice as often as the rewarded ones, the chance level performance was 33.3%.

When the rats reliably responded to the rewarded stimuli (60% or more correct presses for at least four out of five previous training sessions), we gradually shortened the presentation of the stimuli from 90 s to 15 s, and we changed the schedule of reinforcement from continuous to the variable ratio 3 in the final training configuration. Therefore, both rewarded stimuli were displayed 18 times, and both non-rewarded stimuli 36 times during the final training sessions. Blank pauses between stimuli lasted 3 s, and the final training session lasted 32 min 24 s.

The pseudorandom sequences of particular stimuli repeated during the training sessions of the dynamic and static versions of the VSD task are described below. These sequences were repeated six times during a single final training configuration session.

VSD task – dynamic version, first apparatus - training sessions: Static REW, Static non-REW, Dynamic REW, Dynamic non-REW, Static non-REW, Dynamic REW, Dynamic non-REW, Static non-REW, Dynamic non-REW, Static REW, Dynamic non-REW, Dynamic REW, Static non-REW, Dynamic non-REW, Static REW, Static non-REW, Dynamic non-REW, Static non-REW.

VSD task – dynamic version, second apparatus - training sessions: Static REW, Static non-REW, Dynamic non-REW, Static non-REW, Dynamic REW, Dynamic non-REW, Static non-REW, Dynamic REW, Dynamic non-REW, Static REW, Dynamic non-REW, Static non-REW, Dynamic non-REW, Dynamic REW, Static non-REW, Dynamic non-REW, Static REW, Static non-REW.

VSD task – static version, first apparatus - training sessions: Static REW 1, Static non-REW 1, Static REW 2, Static non-REW 2, Static non-REW 1, Static REW 2, Static non-REW 2, Static non-REW 1, Static non-REW 2, Static REW 1, Static non-REW 2, Static REW 2, Static non-REW 1, Static non-REW 2, Static REW 1, Static non-REW 1, Static non-REW 2, Static non-REW 1.

VSD task – static version, second apparatus - training sessions: Static REW 1, Static non-REW 1, Static non-REW 2, Static non-REW 1, Static REW 2, Static non-REW 2, Static non-REW 1, Static REW 2, Static non-REW 2, Static REW 1, Static non-REW 2, Static non-REW 1, Static non-REW 2, Static REW 2, Static non-REW 1, Static non-REW 2, Static REW 1, Static non-REW 1.

**VSD task - dynamic version: Generalization test**

During this generalization test, we displayed the four novel non-rewarded dynamic stimuli pseudorandomly between the presentations of the four familiar training stimuli. Each novel dynamic stimulus (Compl. REW, Compl. non-REW, Control 1, Control 2) was displayed 12 times during the generalization test (six consecutive repetitions of the pseudorandom sequence of stimuli – see below). Individual presentations lasted only 10 s to decrease the likelihood of learning that the novel stimuli are non-rewarded. Single dynamic circle approaches lasted 1 s, and the circles moved at 12.45 cm/s. Therefore, the complete novel dynamic stimuli consisted of 10 repetitions of a particular approach. Both familiar rewarded stimuli (Static REW, Dynamic REW) were displayed 18 times, and both familiar non-rewarded stimuli (Static non-REW, non-Dynamic REW) were displayed 36 times during the generalization test. The blank pauses between individual stimuli lasted 3 or 5 s (pseudorandom distribution), and one complete generalization test took 45 min 24 s. The pseudorandom sequence of stimuli was the same for both subgroups. This was done in order to eliminate any possible bias due to the pseudorandom arrangement of stimuli. The sequence is described below and was repeated six times during the generalization test.

VSD task – dynamic version, both apparatuses - generalization tests: Static REW, Dynamic non-REW, Static non-REW, Compl. REW, Dynamic non-REW, Compl. non-REW, Dynamic REW, Static non-REW, Static REW, Static non-REW, Control 2, Dynamic non-REW, Control 1, Static non-REW, Dynamic REW, Dynamic non-REW, Control 2, Static non-REW, Compl. non-REW, Dynamic REW, Dynamic non-REW, Dynamic non-REW, Compl. REW, Static REW, Static non-REW, Control 1.

**VSD task - static version: Generalization test**

The experimental design of this generalization test was the same as for the VSD task - dynamic version: Generalization test except for the set of used stimuli. In the novel dynamic stimuli, the upper/lower circles were moving at 6.23 cm/s, and the circles that started in the center of the screen had a speed of 8.8 cm/s. The pseudorandom sequence of stimuli displayed during this generalization test is described below and it was repeated six times during a single test session.

VSD task – static version, both apparatuses - generalization tests: Static REW 1, Static non-REW 2, Static non-REW 1, Compl. REW 1, Static non-REW 2, Compl. non-REW 1, Static REW 2, Static non-REW 1, Static REW 1, Static non-REW 1, Compl. non-REW 2, Static non-REW 2, Compl. REW 2, Static non-REW 1, Static REW 2, Static non-REW 2, Compl. non-REW 2, Static non-REW 1, Compl. non-REW 1, Static REW 2, Static non-REW 2, Static non-REW 2, Compl. REW 1, Static REW 1, Static non-REW 1, Compl. REW 2.

**Data Analysis**

We calculated the mean probability as the average number of lever presses per second within the presentation periods. As this probability was not independent for each animal, data was analyzed using a generalized linear mixed model (GLMMN), taking the rat as a grouping factor (i.e., including a random intercept for the rat). Due to the dichotomous response variable (probability of pressing the lever), we used a binomial distribution with a logit link function (i.e., a logistic regression model). The stimulus type was the only fixed effect in analyzing the mean probability of lever presses during the asymptotic and generalization performance. The exponential value of each regression coefficient obtained from the model was the Odds Ratio (OR) of each stimulus type, i.e., the multiplying factor of the Odds of each stimulus type, which was then transformed into a probability. To study the learning dynamics of each type of stimulus, we also included the training time as a fixed factor and its interaction with the type of stimulus. As individual rats took different amounts of training sessions to reach the learning criteria during the first training configuration, this time was normalized between 0 and 1, with 0 being the first session and 1 being the last session in the first training configuration. Moreover, we included a random slope of time in the model. Time coefficients obtained from the models were interpreted in the same way. Their exponential values correspond to the OR (increase/decrease of the Odds during the entire training configuration) and were transformed into probability at each session/time.

Probability distribution was analyzed similarly, but in this case, the response variable was calculated as the number of lever presses per ms within 200 ms bins over each presentation period. A mixed effect logistic regression (GLMM with binomial distribution and logit link function) was also used, with the stimulus type, the time from the stimulus onset (binned into 200 ms periods), and its square as fixed factors, both in interaction with the stimulus type and with a random intercept with each rat as grouping factor. This analysis was only done to study asymptotic and generalization performance.

We calculated the time from the onset of the stimulus to the first lever press for each presentation period to analyze the reaction time. This kind of response variable (time-to-event) is suitably analyzed with Survival models. In this case, we chose a Cox Proportional Hazards model. The exponential of the coefficients obtained from the model corresponds to the Hazard Ratio (HR). The model calculates an empirical baseline hazard function over the time course of the presentation period (which corresponds to the hazard, i.e., the "risk" or probability of suffering the event, when all independent variables are 0) and assumes that a factor proportionally modifies this hazard over all time points of the presentation period. This way, at any time from the onset of the stimulus, the probability of pressing the lever (the risk in our case) is multiplied by the HR, so HRs greater than 1 implies a greater risk at any time, which in turn implies that it is more feasible that the rat presses the lever for the first time earlier, while HRs lower than 1 implies a decrease of the risk, so rats would tend to press the lever later. For analyses of asymptotic and generalization performance, we included the stimulus type as a fixed effect factor, and to avoid the lack of independence within rats, a random intercept was also included (taking the rat as a grouping factor). Data were plotted as cumulative hazard functions representing the probability that the rat has already pressed the lever for the first time (y-axis) at a certain time from the onset of the stimulus (x-axis). The faster the curve grows, the greater the HR, and the earlier the rat can press the lever for the first time. Median lifetime was also obtained, representing the median reaction time of the rats. A normalized time (as in above) was included as a fixed factor to get the HR for each stimulus type at each session of the training configuration to analyze learning dynamics. The HR of each stimulus type was referred to as its baseline risk, i.e., the risk of each stimulus type in the first session, so the probability or risk that appears in the y-axis is an arbitrary value (as risk is the function of the reaction time). Information about the dynamics is extracted from the curves' shapes, so curves that grow faster imply that the reaction time decreases faster than those that grow slower.

### SUPPLEMENTAL DISCUSSION

#### The novelty effect of stimuli in the generalization tests of the dynamic version of the VSD task

The rats' lever pressing probability was lower during the presentation of the novel Compl. non-REW stimulus, which shared the position of predicted collision with the Dynamic non-REW stimulus from the training phase, compared to the novel Control 1 stimulus, but not the novel Control 2 stimulus (Fig. 3D-3E).

This discrepancy might be caused by the difference in the design of the two novel control stimuli. Control 1 stimulus had the starting positions of the moving circles identical to the locations of the familiar Static REW and Static non-REW stimulus, respectively. Therefore, the rats could perceive the Control 1 stimulus as potentially rewarding because the starting position of one circle was previously rewarded. However, the starting position of the other circle was identical to the previously non-rewarded static position. However, there was no difference in the probability of lever pressing between the two novel control stimuli (Fig. 3D-3E). Altogether, these results suggest that the objects' starting positions in our task do not present the crucial information used to generalize novel stimuli, but rather, they are the predicted intersection of the displayed moving objects.

During individual presentations of the novel stimuli in the generalization tests of the VSD task - dynamic version, the lever pressing probability was initially equivalent for all four novel stimuli but changed over their duration. It gradually increased for the Compl. REW stimulus and progressively decreased for the Compl. non-REW stimulus, respectively. At the same time, it stayed similar for the whole duration of both control stimuli (Fig. 3E). The equal probability of lever pressing for all four novel stimuli within the first few seconds of their presentations complies with the natural tendency of rats to respond to novel stimuli[1,2]. However, after the first two 1-s repetitions of circle approaches during novel stimuli, when the rats could estimate the predicted collision position more reliably, they clearly generalized the Compl. REW and Compl. non-REW stimuli. These results support the previously mentioned interpretation that the position of the predicted collision is crucial for discriminating and generalizing the dynamic visuospatial stimuli in our task.

### SUPPLEMENTAL FIGURES AND TABLES

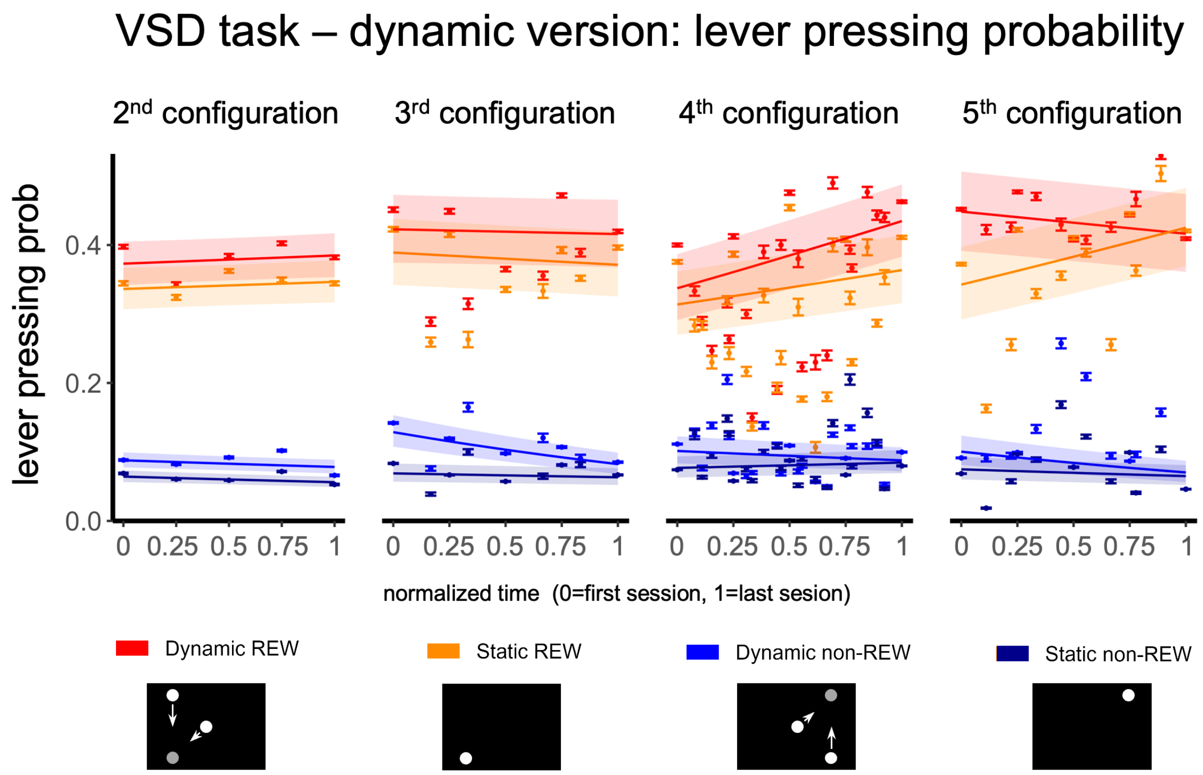


**Supplemental Figure 1**: Mean probability of pressing the lever within 1 s bins of the presentations of the four types of familiar stimuli during the 2nd-5th configuration (from left to right) of the training phase of the dynamic VSD task. Points represent mean ± SEM. Lines represent predictions from logistic regression (GLMM with binomial distribution and logit link function with grouping variable the rat id. Fixed factors: stimulus type, normalized time, and their interaction).

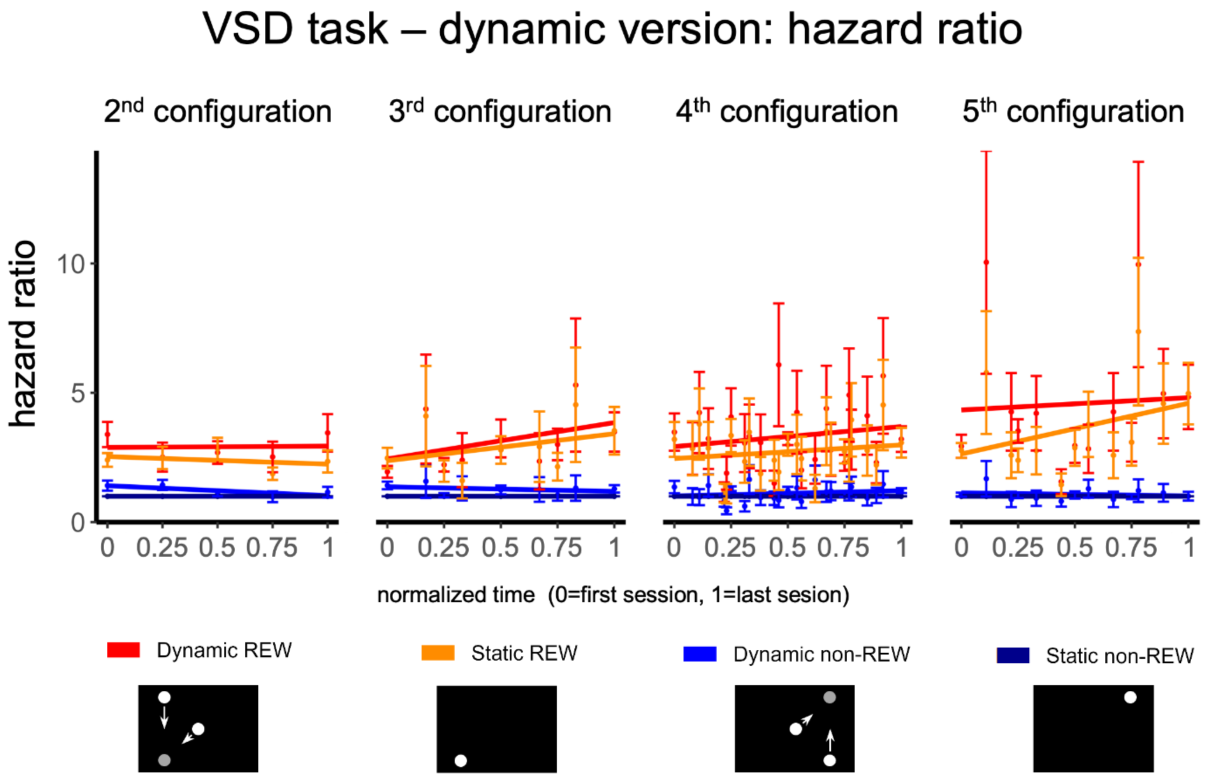


**Supplemental Figure 2**: Hazard Ratio (HR) for the four types of familiar stimuli during the 2nd-5th configuration (from left to right) of the training phase of the dynamic VSD task. Points represent HR ± SEM from Cox Proportional Hazards regression models at different training times (grouping variable the rat id and stimulus type as a fixed factor). Lines represent HR trends with normalized time from linear regression (only for illustrative purposes).

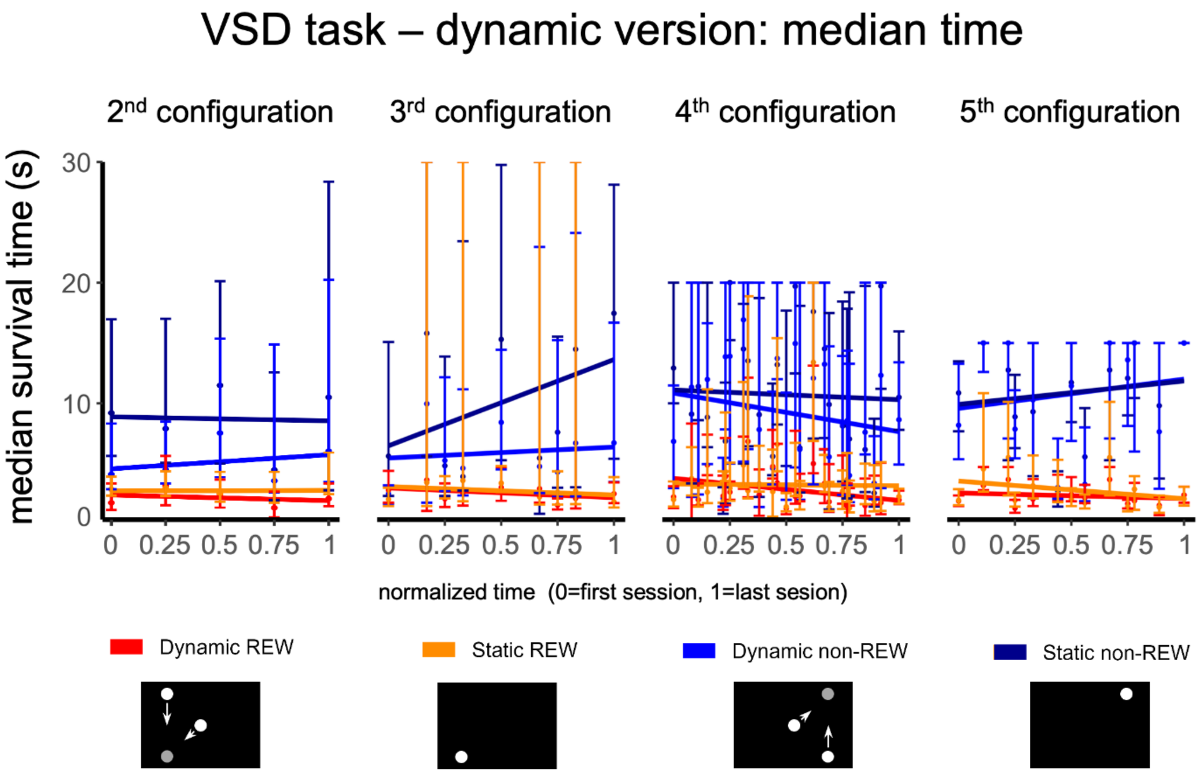


**Supplemental Figure 3**: Median survival time (i.e., the median reaction time of pressing the lever for the first time after the stimulus onset) for the four types of familiar stimuli during the 2nd-5th configuration (from left to right) of the training phase of the dynamic VSD task. Points represent Median Survival Time ± SEM for each stimulus type from survival curves at different training times. Median survival time means the shortest survival time for which the survivor function is less than or equal to 0.5. Lines represent trends of Median Survival Time with normalized time from linear regression (only for illustrative purposes).

| Configuration | Time/stimulus | Reinforcement | Session duration | Pause | S REW | S non-REW | S GEN |
|---|---|---|---|---|---|---|---|
| 1st | 90 s | CR | 28 min 30 s | 5 s | 3 | 6 | --- |
| 2nd | 45 s | FR2 | 30 min | 5 s | 6 | 12 | --- |
| 3rd | 30 s | VR3 | 30 min | 3 s | 9 | 18 | --- |
| 4th | 20 s | VR3 | 34 min 30 s | 3 s | 15 | 30 | --- |
| 5th | 15 s | VR3 | 32 min 24 s | 3 s | 18 | 36 | --- |
| Gen | 15 + 10 s | VR3* | 45 min 24 s | 3 + 5 s | 18 | 36 | 12 |

**Supplemental Table 1**: The design of the behavioral configurations used for training in the VSD task. Configurations 1st – 5th were the training configurations, and Gen was the configuration for generalization tests. Time/stimulus shows the duration of a single stimulus in the individual configuration. Reinforcement: CR = continuous reinforcement, the reward was delivered after every correct lever press; FR2 = fixed ratio 2, the reward was delivered after every second correct lever press; VR3 = variable ratio 3, the reward was delivered after three presses on average but never after more than five correct presses. Session duration shows the total time to complete a single session. Pauses refer to the blank screen periods between individual stimuli presentations. For the generalization test, 3- and 5-s pauses were pseudorandomly presented between individual stimuli. S REW = the number of presentations of each type of rewarded stimulus during the session, S non-REW = the number of presentations of each type of non-rewarded stimulus during the session, and S GEN = the number of presentations of each novel stimulus during the generalization test. *novel stimuli in the generalization test were not rewarded.

| Animal | 1st config. | 2nd config. | 3rd config. | 4th config. | 5th config. | Total sessions |
|---|---|---|---|---|---|---|
| rat 39 | 17 | 5 | 5 | 10 | 10 | 47 |
| rat 41 | 22 | 5 | 7 | 5 | 5 | 44 |
| rat 42 | 11 | 5 | 5 | 14 | 5 | 40 |
| rat 43 | 31 | 5 | 5 | 6 | 11 | 58 |
| rat 44 | 30 | 5 | 5 | 5 | 5 | 50 |
| rat 45 | 22 | 5 | 5 | 5 | 5 | 42 |
| rat 46 | 21 | 5 | 5 | 5 | 5 | 41 |

**Supplemental Table 2**: The learning progress in the dynamic version of the VSD task. The number of sessions to achieve stable performance in the respective configuration of the task and the total number of training sessions for each rat to reach the criteria in the final (5th) training configuration. Rat 40 was excluded from the training after 45 sessions as it could not reach a stable performance of > 60% in the first configuration of the task.
